# Wine terroir and the soil microbiome: an amplicon sequencing–based assessment of the Barossa Valley and its sub-regions

**DOI:** 10.1101/2020.08.12.246447

**Authors:** Jia Zhou, Timothy R. Cavagnaro, Roberta De Bei, Tiffanie M. Nelson, John R. Stephen, Andrew Metcalfe, Matthew Gilliham, James Breen, Cassandra Collins, Carlos M. Rodríguez López

## Abstract

Soil is an important factor that contributes to the uniqueness of a wine produced by vines grown in specific conditions. Recent data shows that the composition, diversity and function of soil microbial communities may play important roles in determining wine quality and indirectly affect its economic value. Here, we evaluated the impact of environmental variables on the soil microbiomes of 22 Barossa Valley vineyard sites based on the 16S rRNA gene hypervariable region 4. In this study, we report that environmental heterogeneity (soil plant-available P content, elevation, rainfall, temperature, spacing between row and spacing between vine) caused more microbial dissimilarity than geographic distance. Vineyards located in cooler and wetter regions showed lower beta diversity and a higher ratio of dominant taxa. Differences in microbial community composition were significantly associated with differences in fruit traits and in wine chemical and metabolomic profiles, highlighting the potential influence of microbial communities on the phenotype of grapevines. Our results suggest that environmental factors affect wine terroir, and this may be mediated by changes in microbial structure, thus providing a basic understanding of how growing conditions affect interactions between plants and their soil microbiomes.

## 1 Introduction

Wine price differs considerably depending on its quality (e.g., flavor, color and typicity), which is largely determined by the interactions between the grape and the growing conditions including climate, soil, topography, agricultural management, and the wine making process (Bokulich et al., 2016). These interactions influence the expression of wine’s terroir (Bokulich et al., 2016; Jullian Fabres et al., 2017). Research on the drivers of terroir have predominantly focused on abiotic environmental factors, such as climate, soil, viticultural management and wine making process, studied individually (Mira de Orduña, 2010; Romero et al., 2016; Vega-Avila et al., 2015) and simultaneously (Van Leeuwen et al., 2004). However, little research has been done, in the context of terroir, on whether soil microbiomes exhibit distinct patterns of distribution at small geographic scales (e.g. neighboring vineyards), and whether vineyard microbiomes are associated with a wine’s terroir (Burns et al., 2015; Bokulich et al., 2016).

Soil microbiomes, especially bacterial species, have been found to be qualitatively and quantitatively different between vineyard systems (Vega-Avila et al., 2015). Environmental factors, such as topography, climate, soil properties, cultivars and agricultural management, combine to affect soil microbial communities (Reeve et al., 2010; Castro et al., 2010; Lamb et al., 2011). It has been shown that climate and topography, including rainfall pattern and temperature, affect these communities through their impacts on soil (Burns et al., 2015). Soil properties such as soil texture, nitrogen (N) content, phosphorus (P) content, carbon to nitrogen (C:N) ratio, water content, and pH show significant effects on the diversity and composition of microbial communities (Girvan et al., 2003; Frey et al., 2004; Rousk et al., 2010; Fierer and Jackson, 2006). Plant genotypes exert an influence on the structural and functional diversity of soil microbiomes by varying root exudates and rhizodeposition (Broeckling et al., 2008; Dias et al., 2013; Philippot et al., 2013). Management practices, land use and varying degrees of stress and disturbance influence the soil microbiome markedly due to specific management objectives (Crowder et al., 2010; Lumini et al., 2011; Reeve et al., 2010; Sugiyama et al., 2010).

Soil microbiomes interact with the vines, and thus affect wine quality (Bokulich et al., 2016; Burns et al., 2015). The interaction between soil microorganisms and plants includes the facilitation of nutrient uptake/utilization, stabilization of soil structure, reduction of disease prevalence by out-competing soil-borne pathogens or increase of disease prevalence by microbial pathogen invasion (Edwards et al., 2014; Zarraonaindia et al., 2015). Soil microbiomes also contribute to the wine fermentation flora, ultimately affecting wine quality (Barata et al., 2012; Compant et al., 2011; Martins et al., 2013). However, microbial assemblage function is intrinsically difficult to measure and define because of its highly changeable nature (Nannipieri et al., 2003). Additionally, due to the complex interactions between soil microbes, the influence of certain microbial communities can be substituted by other microorganisms with the same ecological function (Nannipieri et al., 2003; Crowder et al., 2010; Lamb et al., 2011; Wittebolle et al., 2009).

The primary aim of this project was to assess if there is a relationship between soil microbiomes and terroir. To achieve this, we asked the following questions:

i. Do wine sub-regions have distinct soil bacterial communities?;
ii. What environmental conditions and agricultural practices shape soil bacterial community of vineyards?; and
iii. Do differences in the soil bacterial community correlate with berry and wine characteristics?

In order to answer these questions, we undertook a soil microbiome survey in an iconic wine region, the Barossa in South Australia. The Barossa has a winemaking history of over 160 years and because of its importance as a growing region, has been chosen as a model to investigate terroir previously (Wolf et al., 2003; Edwards et al., 2014; Xie et al., 2017). Besides, the environmental characteristics of the Barossa, including climate, soil and topography have been previously characterized in detail (Robinson and Sandercock, 2014). However, to date, no study has analyzed the soil microbiomes of the Barossa wine region or the possible influence on wine properties. Thus, determining how soil microbiome diversity and composition are influenced by environmental factors, and how microbiome differences correlate with differences in fruit/wine composition, will provide a starting point from which to better understand the (potential) functional role of soil microbial communities in terroir.

## 2 Materials and methods

### 2.1 Experimental design and plant material

Twenty-two Barossa vineyards (Figure S1), planted with own-rooted Shiraz (*Vitis vinifera* L.) and representative of the climate, soil and management practices of six Barossa sub-regions (i.e. Eden Valley, Northern Grounds, Central Grounds, Eastern Edge, Western Ridge, Southern Grounds) were selected for this study. Three to four vineyards per sub-region were included and nine vines from three rows from each vineyard were selected for measurement and sampling. Vines within the same row were adjacent to each other. Vines adjacent to missing vines, end of row vines and border rows were excluded from the selection.

### 2.2 Soil sampling protocol

Three soil cores (0-10cm soil layer) were collected using a (20 mm diameter) soil auger from around each individual plant (approximately 10cm from the trunk) and combined, giving a total nine soil samples per row. A total of 594 soil samples were collected (27 soil samples from each vineyard) on the 2^nd^ of November (Austral Spring) 2015. Soil samples were immediately stored at 4°C and returned to the laboratory on the same day of collection. Soil samples from the same row were thoroughly mixed to obtain three samples per vineyard, and a total of 66 samples across the study. Coarse debris was removed from each soil sample using a 2mm sieve, and each sample was then divided into three sub-samples (approximately 850 cm3 each). The first subsample (approx. 20 g) was used for determination of soil gravimetric moisture content. The second was air-dried until a constant mass was achieved and used for analysis of soil texture, pH, electrical conductivity, and plant-available (Colwell) P (phosphorus), as described previously (Cavagnaro, 2015)). The third soil subsample was stored at - 80°C for DNA extraction and downstream genomic analysis (see below).

### 2.3 Vineyard physical characterization

In this study, the climate was characterized on the basis of rainfall and temperature. The influence of topography was studied through elevation above sea level and vineyard orientation. Soil texture was determined following (Giddings, 2015). Soil pH and electrical conductivity were determined on a 1:5 soil/water mixture and then measured using pH/salinity meter (WP-81 Conductivity-Salinity-pH-mV Meter, v6.0, TPS Pty Ltd). Plant-available phosphorus was extracted and measured using Colwell P method (Rayment and Higginson, 1992) (Table S1). The remaining soil, topographic and climatic data was obtained from the Barossa Grounds project (Robinson and Sandercock, 2014), while vineyard management information was collected from participating growers (Table S2).

### 2.4 Fruit and wine chemical analysis

Fruit juice pH and total acidity (TA) was measured using an autotitrator (Crison instruments Barcelona, Spain) (Iland et al. 2004). Total soluble solids (TSS) of juice samples were tested with a digital refractometer (BRX-242 Erma inc. Tokyo, Japan). A sample of 50 berries from random bunches on selected vines were collected and frozen at −20°C for anthocyanin, phenolic and tannin analyses. Total grape tannins were measured by the methyl cellulose precipitable (MCP) tannin assay (Sarneckis et al. 2006) using the protocol of Mercurio et al. (2007). Total anthocyanin and phenolics were determined according the method of Iland et al. (2004) (Table S3).

One bottle of commercial wine per vineyard was used for the chemical analysis. Wine pH and TA was determined as described by Iland et al. (2004). Final alcohol levels were determined using an Alcolyzer Wine ME (Anton Paar, Graz, Austria). Wine colour was determined using the modified Somers assay using a high throughput method in 96 well plates [98]. Wine tannin concentration was determined using the methyl cellulose precipitable (MCP) tannin assay of Mercurio et al. (2007) and is expressed as epicatechin equivalents (mg/L) using an 8-point epicatechin standard curve Sarneckis et al. (2006). The modified Somers assay was used to determine; wine colour density (WCD), SO2-corrected WCD, degree of anthocyanin ionisation, phenolic substances and anthocyanins (in mg/L) (Table S4).

Non-targeted metabolomic analysis of the wine samples was performed using LC-MS/MS. The metabolites were isolated from bottled wine samples using solid-phase extraction (SPE) with Phenomenex Strata-X 33 um 85Å polymeric reverse-phase 60mg/3mL cartridges. A 2 mL aliquot of each sample was evaporated to dryness under nitrogen at 30°C. SPE conditions are presented in Table S5. A pooled mix of all samples was prepared and used to monitor instrument performance. The analysis was performed on an Agilent 1200SL HPLC coupled to a Bruker microTOF-Q II in ESI negative mode. The operating conditions are described in Table S5 (Table S6-7).

Following data acquisition, mass calibration was performed on each file using Bruker Daltonic’s DataAnalysisViewer4.1 “Enhanced Quadratic” calibration method (Bruker Singapore, The Helios, Singapore). Each file was exported from DataAnalysis in the mzXML generic file format for further processing. The files were processed using R (statistical programming environment) v3.1.0 and Bioconductor v2.14 under a Debian Linux 64-bot environment. Molecular features were extracted for each file using xcmx package and features that possessed a common mass and retention time across samples were grouped together.

### 2.5 16S rRNA Gene Next Generation Sequencing library preparation

DNA extractions from soil 66 samples were carried out at the Australian Genome Research Facility (AGRF) (Adelaide node) using MoBio Powersoil kit (MoBio Laboratories, Inc) following the manufacture’s protocol. DNA concentrations were estimated using a Nanodrop 2000 spectrophotometer (Thermo Fisher Scientific, Waltham, MA, USA) and normalized to 5 ng/μl using nanopure water.

Primers 515F and 808R (Bates et al., 2011; Caporaso et al., 2011) specific for the Bacterial 16S rRNA gene hypervariable “V4” region were used for PCR amplification of extracted DNA and to prepare amplicon libraries. 515F worked as a universal forward primer for all the samples and 808R included 12-base sample specific barcodes to allow downstream de-multiplexing (Table S8).

Three replicated PCR reactions were performed for each of the 66 samples. Each of these runs included one negative control as ‘sample67’ with no template DNA added. PCR reactions included 10ng of extracted DNA, 12.5 μl Q5 high-fidelity 2*master mix (New England Biolabs), 8.5 μl dH_2_O, 1μl forward and reverse primers (10 μM) in 25μl reaction system. The PCR thermocycler (Bio-Rad T100) program was 95°C for 6min, followed by 38 cycles of 95°C for 30sec, 50°C for 30sec and 72°C for 1m30sec.

Success of PCR reactions was verified by agarose gel (1.5% w/v) electrophoresis. Samples exhibiting weak bands were reamplified by adjusting the amount of DNA template. The triplicate reactions were then pooled into 67 pools. Individual pools were quantified by Qubit fluorometric double stranded DNA assay (Invitrogen, Carlsbad, CA, USA) and then mixed on an equimolar base to generate six pools each with 11 samples (each containing 5 ul of the water control pool). Pools were size-selected to remove unused primers using Agencourt^®^ AMPure^®^ XP (Beckman Coulter, Brea, CA, USA) following the manufacturer’s protocol and mixed to equimolar concentrations to make one final pool. Library concentration and fragment size were estimated using TapeStation (Agilent, Santa Clara, CA, USA) and sequenced on the Illumina MiSeq platform (300 bp PE) (Illumina, San Diego, CA, USA) at the Australian Genome Research Facility-Adelaide node (AGRF).

### 2.6 Bioinformatics analysis

Raw Illumina sequencing data was quality filtered and demultiplexed at AGRF-Adelaide node. Forward and reverse sequences passing QC filter were merged using *bbmerge* (Bushnell, 2016).

Merged reads were analyzed using Quantitative Insight Into Microbial Ecology (QIIME) (QIIME version 1.8.0) (Navas-Molina et al., 2013). Operational taxonomic units (OTUs) were clustered using open-reference picking with the default *uclust* method (Edgar, 2010) based on 97% sequence similarity to the 16S rRNA Greengenes database (McDonald et al., 2012; DeSantis et al., 2006). OTUs were aligned to the Greengenes core reference database using *PyNAST* (Caporaso et al., 2010). Ribosomal Database Project (RDP) classifier was used to assign taxonomy (Wang et al., 2007). Both closed-reference OTU picking and open-reference OTU picking were performed for later analyses.

A BIOM file was generated after OTU picking, then OTUs identified in the negative control samples were removed from soil sample OTUs, leaving between 37,176 and 114,777 OTUs per sample (mean = 60,147 OTUs). Data with and without rarefaction were used for alpha diversity and beta diversity analyses. 37,176 OTUs (the lowest amount of OTUs in one sample) were randomly selected from each sample for rarefaction.

Alpha diversity (within-sample species richness and evenness) was measured using non-phylogenetic (including the observed number of OTUs and the Chao 1 estimator of the total number that would be observed with infinite sampling) and phylogenetic (Faith’s Phylogenetic Diversity) indices (Faith, 1992). Phylogenetic beta diversity (between-sample diversity) was calculated using both weighted and unweighted UniFrac (Lozupone and Knight, 2005) and three-dimensional principal coordinates analysis (PCoA) plots were built through Emperor (Vázquez-Baeza et al., 2013). We then constructed a neighbor joining ultrametric tree in QIIME from the beta diversity UniFrac distance matrix. The generated tree file, as well as the Barossa Valley geographical map, vineyard locations and taxa summary files, were input into GenGIS (Parks et al., 2009, 2013) to visualize the relationship between soil bacterial beta diversity and vineyard location. The statistical significance of this relationship was determined using the Mantel test based on 9,999 random permutations and implemented on GenAlex v6.5 (Peakall and Smouse, 2012).

To identify the association of environmental variables and grape and wine properties (Table S1-S7) with soil bacterial microbiome, bacterial community dissimilarities were visualized with non-metric multidimensional scaling (nMDS) plots. Variables were fitted to the ordination plots using the function *envfit* in the package *Vegan version 2.5-2* (Oksanen et al., 2013) implemented in R version 3.5.0 (Team, 2013). Spearman’s rank correlation coefficients were measured between individual taxon abundance and fruit and wine traits using the function *rcorr* in the package *Hmisc*. Grape traits included those from sensory, basic chemistry analyses, while wine traits included basic chemistry, wine fermentation products and amino acids concentration. Those traits and taxa with a significant (p-value <0.05) correlation coefficient larger than 0.80 or lower than −0.80 were deemed as significantly associated.

To identify which variables are important in explaining the composition of the soil microbial community, we performed distance-based redundancy analysis (dbRDA), a form of multivariate multiple regression that we performed directly on a Bray-Curtis dissimilarity matrix of OTUs using the ADONIS function in *Vegan*. We used automatic model building using the function *step* in R. The step function uses Akaike’s Information Criterion (AIC) in model choice, which is based on the goodness of fit. The model building proceeds by steps until the ‘best’ fit is identified. If two predictor variables were highly correlated (>0.85) one, typically that which was more difficult to measure, was removed as well as variables with missing replicates (Variables included in the automatic model building are marked with * in Tables S2-S8). Differential statistic functions within the *edgeR* package (Chen et al., 2008) was used, as in Weis et al. (2017) to determine the significantly different taxa between vineyards separated by the main environmental drivers of beta diversity (i.e. soil type and soil phosphorous content). In order to avoid the influence of taxa showing low counts, a minimum threshold was set up at 100 counts per million.

## 3 Results

### 3.1 Analysis of soil properties

Of the three soil physicochemical properties tested, plant-available phosphorous (P) and electrical conductivity (a measure of soil salinity), differed significantly (Kruskal-Wallis: p-value < 0.05) between sub-regions of the Barossa (Table S1). Plant-available P was lowest in the Northern Grounds (11.5±2.7μg P/g soil) and highest in the Eastern Edge (39.0 ±14.2μg P/g soil). Electrical conductivity ranged from 111.0 uS/cm (Northern Grounds, SE=34.2) to 302.5 uS/cm (Central Grounds, SE=123.5). Soil pH did not differ between sub-regions, ranging from 6.2 (Eden Valley, SE=0.4) to 6.8 (Southern Grounds, SE=0.5).

### 3.2 Barossa Valley soil bacteria community composition

After quality filtering of the raw sequencing results, an average of 130,949 paired sequences remained per sample. Of these an average of 86,835 paired-end sequences per sample (66.3%) could be joined using *bbmerge* (Table S9).

Both bacterial and archaeal DNA was detected in all soil samples. A total of 98.9% of sequences were classifiable at the phylum level (Figure 1A) and 95.2% at the genus level. Of those classifiable at the phylum level, 96.5% were assigned to one of nine dominant groups (relative abundance ≥1.0%) in the samples namely: Actinobacteria (26.9%), Proteobacteria (26.7%), Acidobacteria (12.0%), Planctomycetes (6.2%), Chloroflexi (5.6%), Firmicutes (5.3%), Gemmatimonadetes (3.9%), Bacteroidetes (3.5%), Verrucomicrobia (2.5%) (Figure 1A). The only dominant Archaea group was Crenarchaeota (4.0%). The overall dominant Bacteria and Archaea groups were consistently present in the six regions, but at different ratios (Figure 1A). The phylogenetic inference of microbiome composition differences between sub-regions showed three clusters with Central and Northern Grounds, and Eden Valley and Western Ridge sharing the more similar microbial profiles (Figure 1A).

**Figure 1.**
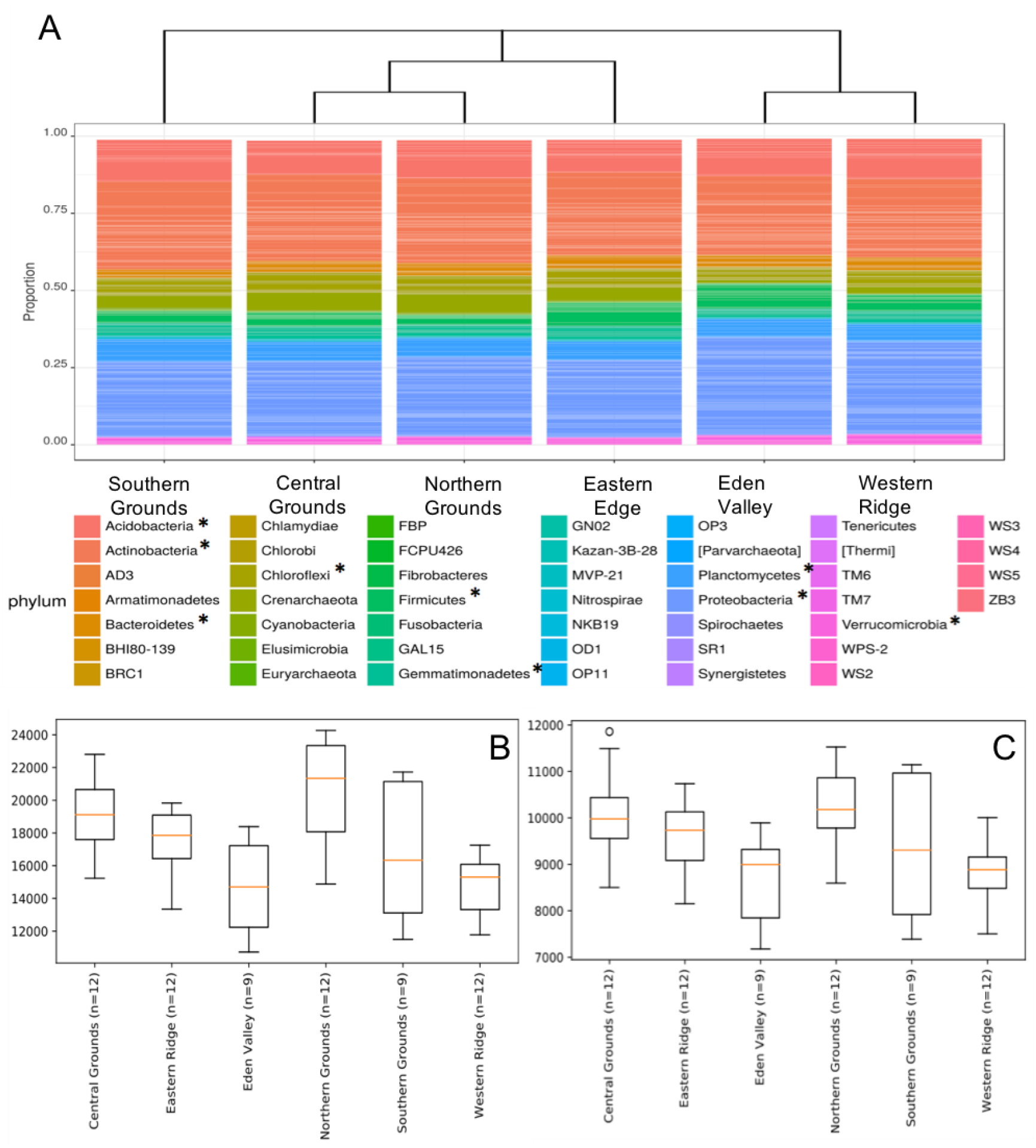
Soil bacteria community composition and diversity in 6 Barossa sub-regions. **A)** Phylogenetic inference of microbiome composition differences between Barossa sub-regions. Neighbour joining tree was generated with weighted UniFrac distances calculated with sequences classifiable at the phylum level (98.9% of total). 96.5% of all sequences were assigned to one of nine main dominant groups (relative abundance >= 1.0%) (indicated here by *). **B)** Alpha diversity: Chao1 diversity comparison, **C)** and observed species diversity comparison. Alpha diversity values were calculated based on rarefied data was established using 16S sequencing reads from 3 soil samples per vineyard.

The number of observed OTUs (Figure 1B) showed significant differences (t-test: p-value < 0.05) between the OTU rich sub-regions (Northern and Central Grounds) and the relatively OTU poor sub-regions (Eden Valley and Western Ridge) (Table S10). Similarly, the Chao1 metric showed that Northern and Central Grounds presented higher levels of OTU richness while Eden Valley and Western Ridge had the lowest (Figure 1C). Pairwise comparison of alpha diversity between sub-regions showed significant differences (t-test, p-value < 0.05) between Northern Grounds and Eden Valley and Western Ridge and between Central Grounds and Eden Valley and Western Ridge (Table S11).

Dissimilarities in microbial communities between samples (i.e. beta diversity) were calculated as weighted and un-weighted UniFrac distances and both methods showed similar patterns, and so only analyses based on weighted results are shown here. For the most part, the three replicates from within a given vineyard were closely grouped on the ordination plot (Figure 2A), indicating that bacterial communities were consistent within sites. Pairwise analysis of the differences between groups (vineyards and sub-regions) showed that all vineyards and sub-regions are significantly different to each other (Adonis, p-value < 0.001). Mantel test analysis of the association between microbiome compositional differences and geographic distance, showed a small but significant correlation (rxy = 0.315; p-value = 0.0001) (Figure 2B).

**Figure 2.**
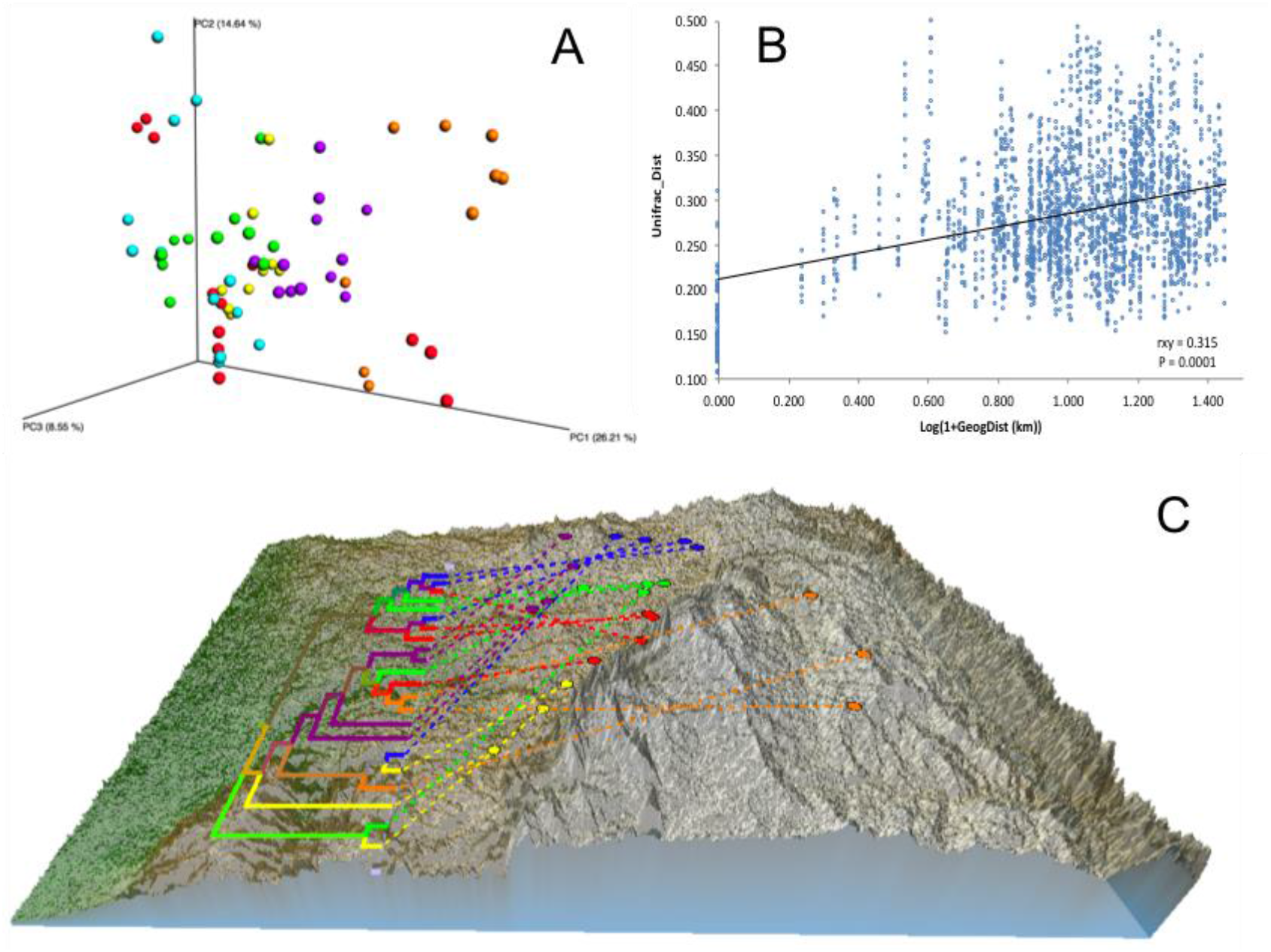
Effect of vineyard location on soil microbiome differentiation. **A)** PCoA based on Beta diversity of soil bacterial communities calculated using weighted UniFrac distances. Values were calculated based on rarefied data to 37,176 sequences per sample. **B)** Relationship between phylogenetic Beta diversity and geographic distance. Unifrac_dist indicates weighted UniFrac distances. Geographic distances were calculated from latitude/longitude coordinates using GenAlex v6.5 *geographic distance* function implemented as Log(1+distances in Kilometres). The relationship was tested using Mantel’s correlation coefficient (rxy) with its probability estimate for significance (P) based on 9,999 random permutations and implemented using GenAlex v6.5. **C)** Neighbour joining ultrametric tree calculated from Beta diversity weighted UniFrac distance matrix between 22 vineyards located in six sub-regions: Northern Grounds (blue); Southern Grounds (yellow); Central Grounds (green); Eastern Edge (red); Western Ridge (purple); Eden Valley (orange). Tree was overlayed with the Barossa Region elevation map using GenGIS. Beta diversity was established using 16S sequencing reads from 3 soil samples per vineyard.

To further explore dissimilarities among and within regions, neighbor joining analysis was used to cluster samples and to generate a similarity tree in QIIME. This information, along with a geographical map of the regions and their locations, were combined using the GenGIS software package (Parks et al., 2009). This approach showed a low level of clustering of vineyards according to their geographic location (Figure 2C).

### 3.3 Drivers of soil microbiome differentiation

Model selection was used to identify the combination of variables that explained the greatest variation in the soil microbiome. This approach consistently selected soil plant-available phosphorus (P) and soil texture as the main drivers (Model: p-value = 0.001) of soil microbiomes in the Barossa vineyards tested (Figure 3). Together, both variables explained 19.7% of the observed variability. Independent pairwise analysis of UniFrac distances of vineyards grouped by these soil characteristics, showed that microbial communities in clay soil types were significantly dissimilar from those in sandy soils (PERMANOVA: p-value < 0.001, Figure 4A). Microbial communities in soils with high plant-available phosphorus (P > 30mg/kg) were also dissimilar from those with low plant available phosphorous (PERMANOVA: p-value < 0.001, Figure 4C). Three and eight taxa were significantly more abundant in clay and sandy soils respectively (Figure 4B), while eight taxa were found significantly associated with low plant available phosphorous content, and three associated high levels of plant available phosphorous in soil (Figure 4D).

**Figure 3.**
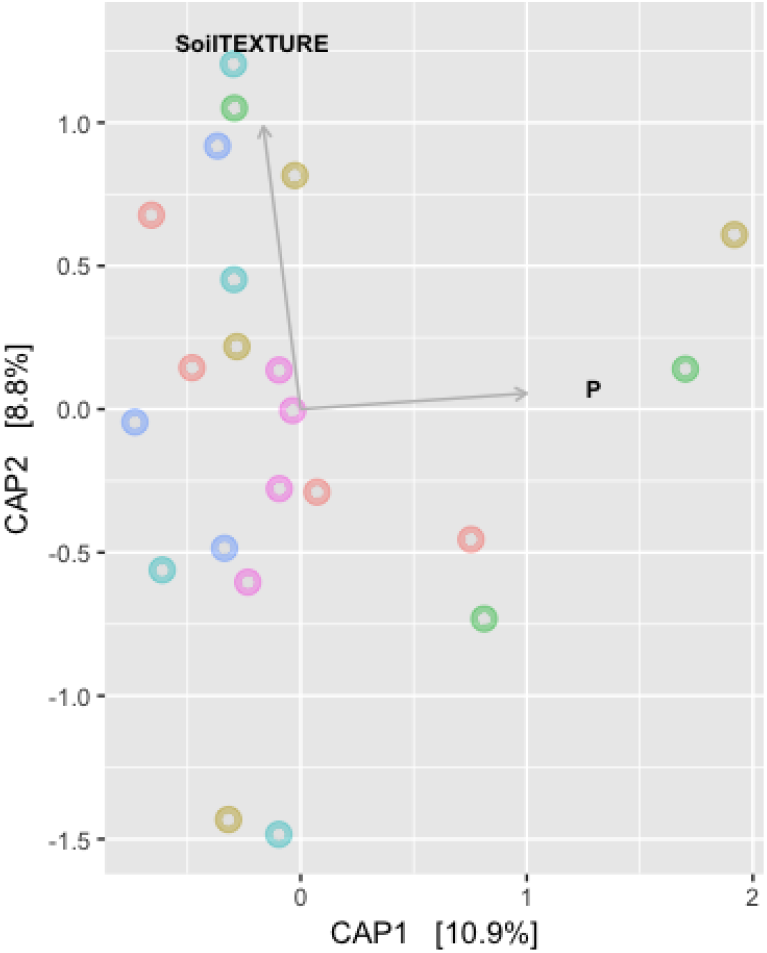
Main drivers of soil microbiome differentiation between Barossa Region vineyards. The observed important soil factors that affect soil microbial community groups in combinations. CAP plot displays the combination of variables that explained the greatest variation in the soil microbiome through model selection (full results Table 1). The correlation test was carried out on environmental variables following the removal of the highly correlated variables (>0.85) using the function *ordisten*, in the package *Vegan*. The variables implemented in the final model were soil phosphorous and soil texture, which explained 19.7% of variation in the soil microbiome. Distance based redundancy analysis (dbRDA) with Bray-Curtis dissimilarity matrix of OTUs was used to examine the influence of these predictor variables using the function capscale in the package Vegan in R.

**Figure 4.**
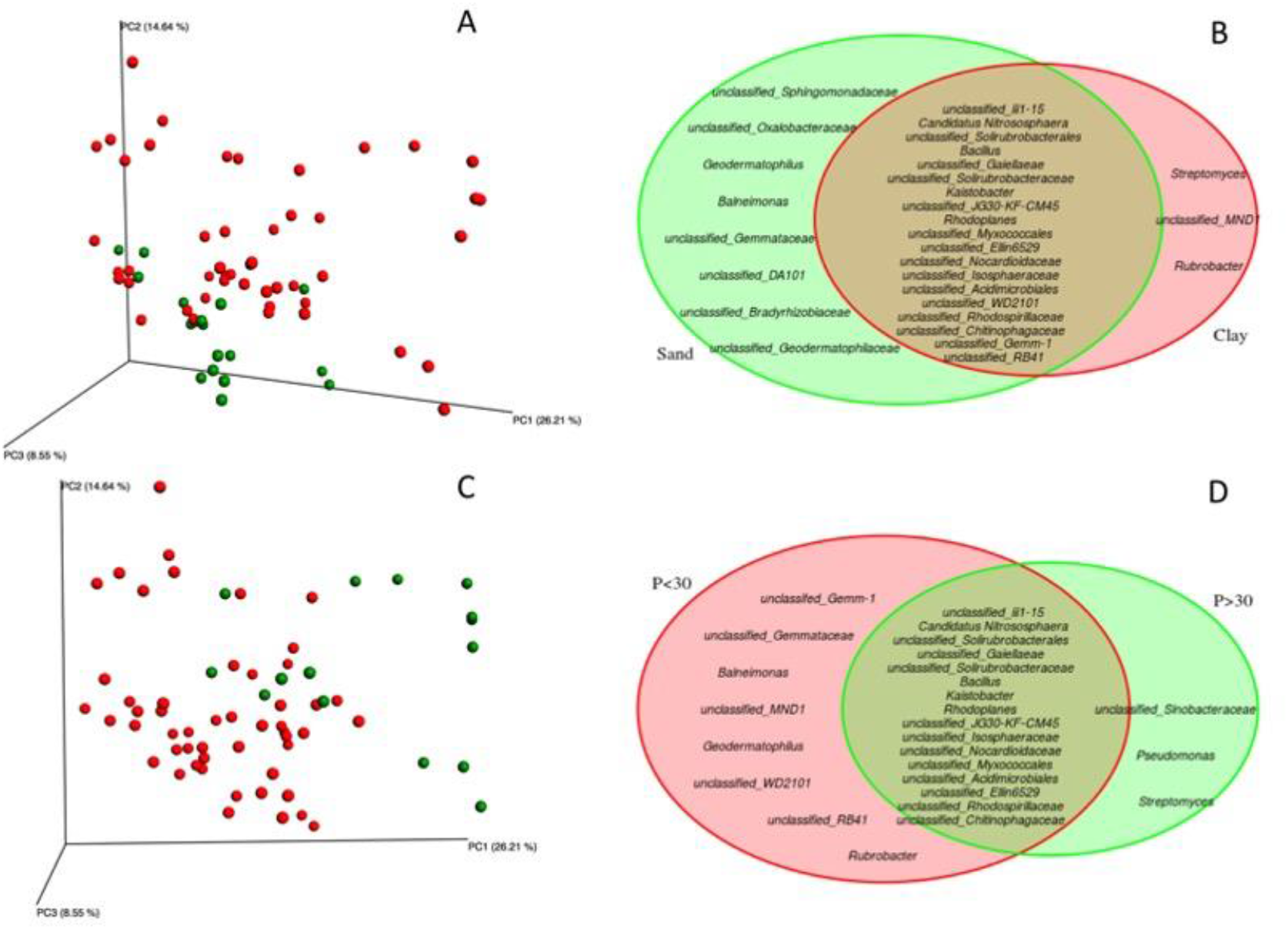
Identification of microbial genera associated to soil texture and plant-available phosphorous in Barossa Region vineyards. Principal coordinate analysis plots display weighted UniFrac distances of soil samples from 22 vineyards in six sub-regions of Barossa Valley. Venn Diagrams show significantly different (P > 0.01) genera. Plots and diagrams are grouped by (A/B) soil type (clay (red) versus sandy soils (green)), and (C/D) plant-available Phosphorous (P) (P < 30 μg P / g soil (red), P > 30 μg P / g soil (green). Beta diversity was established using 16S sequencing reads from 3 soil samples per vineyard.

*Envfit* analysis identified a number of other environmental factors as individually associated with microbial community composition (Figure 5). Aside from plant available phosphorous (r^2^ = 0.3706, p-value < 0.001), these variables were: elevation (r^2^ = 0.3609, p-value < 0.001), growing season rainfall (r^2^ = 0.2499, p-value < 0.001), mean annual rainfall (r^2^ = 0.1621, p-value = 0.004), spacing between rows (r^2^ = 0.1512, p-value = 0.006) and between vines (r^2^ = 0.1561, p-value = 0.011), and growing season mean temperature (r^2^ = 0.1113, p-value = 0.022).

**Figure 5.**
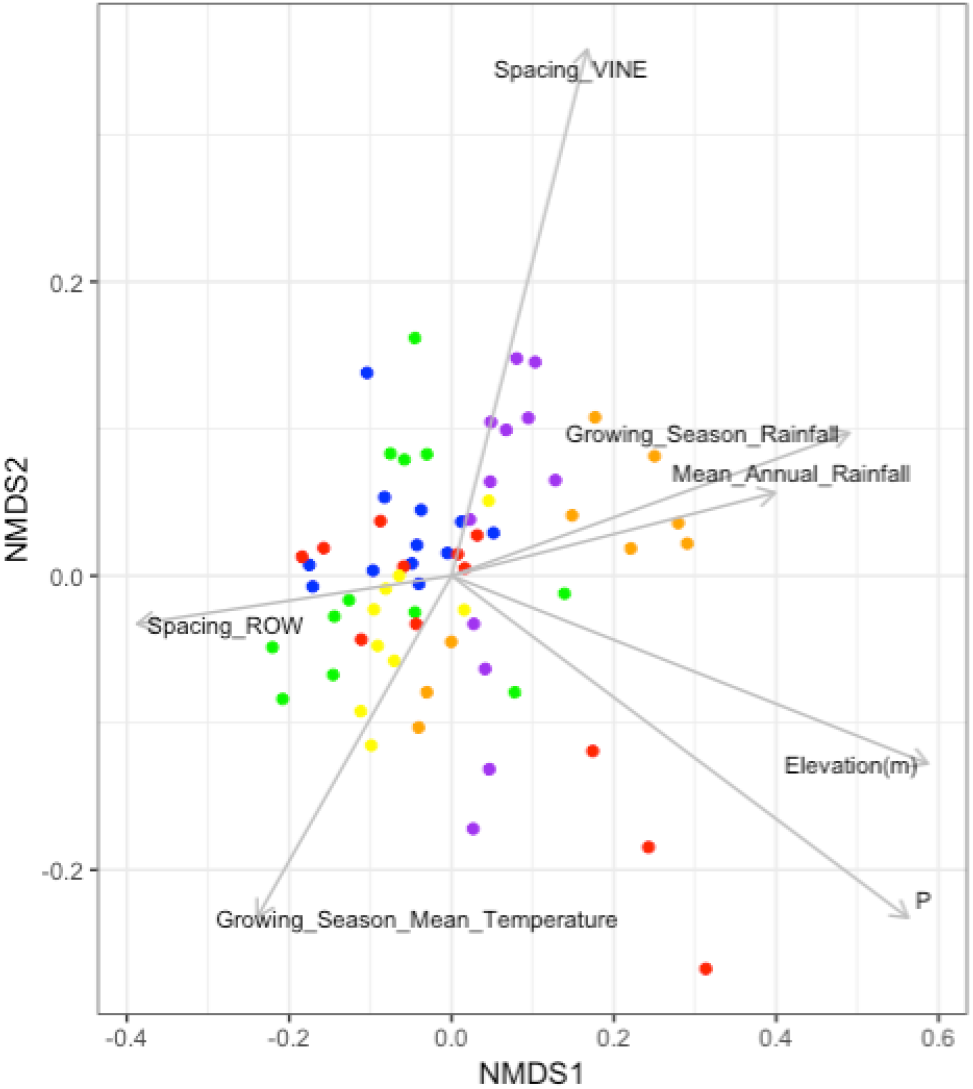
Environmental and vineyard management factors significantly associated with soil microbial community composition in Barossa Region vineyards. Non-metric multidimensional scaling plot displays the microbial community composition of 22 vineyards located in six sub-regions: Northern Grounds (blue); Southern Grounds (yellow); Central Grounds (green); Eastern Edge (red); Western Ridge (purple); Eden Valley (orange). Vector arrows indicate the association with environmental variables with p-value < 0.05. Arrow heads indicate the direction and length indicates the strength of the variable and nMDS correlation. Analysis was conducted using 999 permutations with variables deemed significant where p-value < 0.05.

Analysis of the correlation between individual environmental and vineyard management variables and taxa abundance, identified 4 positive (Spearman’s >0.80, p-value <0.001) and 3 negative (Spearman’s <-0.80, p-value <0.001) significant correlations (Figure S2). Positive correlations with individual taxa included, pH (order iii1-15 and family Pirellulaceae), elevation (family Isosphaeraceae), and plant age (family Hyphomicrobiaceae); while negative correlations included P (family OPB35), elevation (family Conexibacteraceae), and the spacing between vines on the same row (family Haliangiaceae).

### 3.4 Terroir and vineyard soil microbiomes

Twenty four of the 75 grape and wine characteristics included in the study displayed a significant correlation with the soil microbial community composition (Table 2). The strongest associations identified for each of the four groups of traits tested were: 50 berry weight and average color per berry (basic berry properties); total anthocyanins and total phenolics (basic wine chemistry); Glycine and Alanine (wine amino acids); and 2-phenyl ethyl ethanol and acetic acid (wine fermentation products).

**Table 1.**
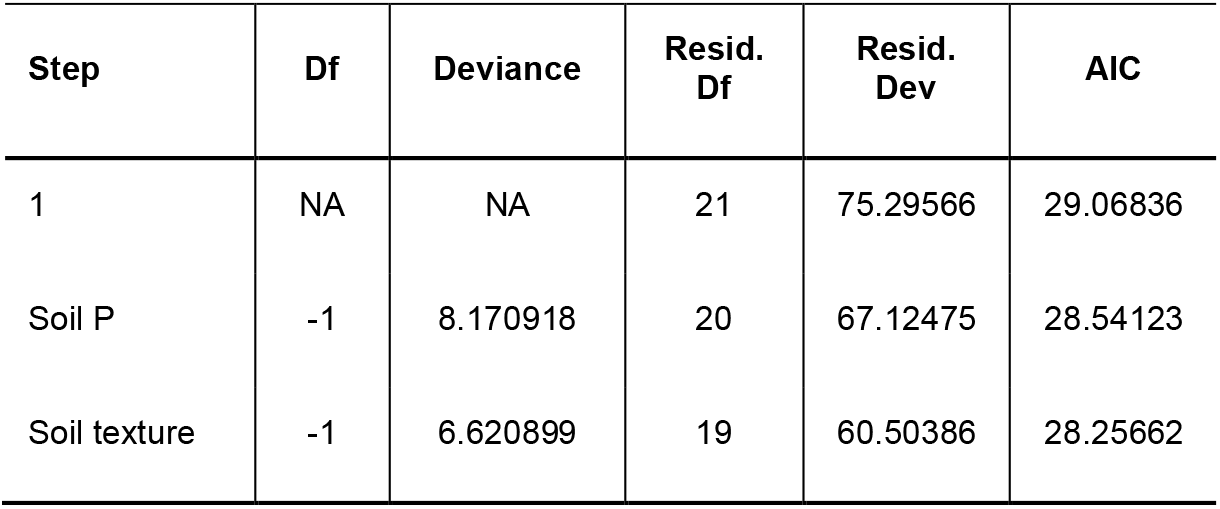
Main drivers of soil microbiome differentiation between Barossa Region vineyards. Variables that explained the greatest variation in the soil microbiome through model selection. The correlation test was carried out on environmental variables following the removal of the highly correlated variables (>0.85) using the function *ordisten*, in the package *Vegan*.

| Step | Df | Deviance | Resid.<br>Df | Resid.<br>Dev | AIC |
| --- | --- | --- | --- | --- | --- |
| 1 | NA | NA | 21 | 75.29566 | 29.06836 |
| Soil P | -1 | 8.170918 | 20 | 67.12475 | 28.54123 |
| Soil texture | -1 | 6.620899 | 19 | 60.50386 | 28.25662 |

**Table 2.** Fruit and wine characteristics significantly associated with microbial community composition in Barossa Region vineyards. Table shows the *envfit* output that was carried out the correlation test between grape and wine characteristics variables that fitted onto an ordination of nonmetric multidimensional scaling (nMDS) plots of microbial community data from soils in 22 vineyard sites. Analysis was conducted using 999 permutations with variables deemed significant where p-value < 0.05.

| Variables |  | NMDS1 | NMDS2 | r2 | Pr(>r) |
| --- | --- | --- | --- | --- | --- |
| **Basic berry properties** | 50 berries weight | -0.87544 | -0.48332 | 0.1612 | 0.008** |
|  | TA berry | 0.9369 | 0.3496 | 0.1119 | 0.029* |
|  | Average colour | 0.76859 | 0.63974 | 0.1337 | 0.008** |
|  | Average total phenolics berry | 0.76558 | 0.64334 | 0.135 | 0.015* |
|  | Malic acid | -0.90493 | 0.42557 | 0.104 | 0.03* |
| **Basic wine chemistry** | Total phenolics | **0.83761** | **0.54627** | **0.2132** | **0.002\*\*** |
|  | Total anthocyanins | 0.99519 | 0.09801 | 0.2507 | 0.001*** |
|  | Colour density (so2 corrected) | 0.72894 | 0.68457 | 0.1449 | 0.006** |
|  | Hue | -0.78985 | 0.61331 | 0.1314 | 0.011* |
| **Wine amino acids** | Alanine | **0.11831** | **0.99298** | **0.124** | **0.017\*** |
|  | Asparagine | 0.55124 | 0.83435 | 0.1156 | 0.023* |
|  | Glutamate | 0.39571 | 0.91837 | 0.103 | 0.031* |
|  | Glycine | 0.62968 | 0.77685 | 0.1847 | 0.002** |
|  | Serine | 0.54731 | 0.83693 | 0.0936 | 0.04* |
|  | Threonine | 0.20213 | 0.97936 | 0.0934 | 0.049* |
|  | Tryptophan | 0.56228 | 0.82695 | 0.119 | 0.025* |
| **Wine**<br>**ferment.**<br>**products** | Acetic acid | -0.99914 | -0.04153 | 0.1689 | 0.003** |
|  | Propanoic acid | -0.96827 | 0.24991 | 0.118 | 0.012* |
|  | 3-methylbutanol | 0.99079 | 0.13538 | 0.1184 | 0.02* |
|  | 2-methylbutanol | 0.98968 | 0.14332 | 0.108 | 0.034* |
|  | Butanoic acid | -0.89212 | 0.4518 | 0.1298 | 0.013* |
|  | 2-phenyl ethyl ethanol | 0.70731 | 0.7069 | 0.2064 | 0.001*** |
|  | 2-phenyl ethyl acetate | 0.82725 | 0.56183 | 0.1249 | 0.013* |

Significant positive correlation (Spearman’s >0.80, p-value <0.05) were identified between the abundance of one taxon (order IS_44) and the average level of total phenolics mg/g berry weight (Figure S3A). Similarly, six wine traits showed positive correlations with the abundance of six microbial taxa (Figure S3B-E). Briefly, the genus *Rhodoplanes* was positively associated with the level of wine total phenolics and the family Chitinophagaceae was associated with color density of SO2 corrected wine and with the level SO2 resistant pigments in wine, while the family Kouleothrixaceae was positively correlated with wine color density.

## 4 Discussion

Previous studies have shown that environmental factors (e.g. climate and soil properties) and crop management may affect microbial populations in vineyards (Fierer, 2008; Burns et al., 2015; Weckert, 2016). To better understand how these variables contribute to vineyard microbial communities and how microbial diversity and composition correlate with fruit and wine quality traits, we studied the soil microbiome composition of 22 commercial vineyards representative of the Barossa Valley wine region in Australia.

### 4.1 Vineyard soil microbiome composition and diversity

With over 37,176 sequences per sample we reached a sequencing depth higher than those achieved in previous studies deemed sufficient to resolve differences between similar samples (e.g. Liu et al., 2014; Burns et al. 2015). From a species composition point of view, our results indicate that vineyard soil microbiomes present similarities across the six sub-regions studied. All soils analyzed presented both bacteria and archaea. A total of 96.5% of the all identified sequences were allocated in one of nine main dominant phyla (relative abundance >= 1.0%). Of these, eight (Actinobacteria, Proteobacteria, Acidobacteria, Planctomycetes, Chloroflexi, Firmicutes, Gemmatimonadetes, Bacteroidetes, and Verrucomicrobia) were Eubacteria, while only one dominant taxon was from Archeabacteria (Crenarchaeota). Although dominant phyla were consistently found in the six regions tested, they were present in different ratios. This finding is similar to earlier work; for example, investigating the Napa Valley American Viticultural Area (AVA), Burns et al. (2015) found the same nine top dominant bacteria groups, also with different ratios and order for each group. Similarly, Liu’s et al. (2014) analysis of agricultural black soils in northeast China found almost the same dominant bacterial groups. Equally, analysis of non-agricultural soils by Faoro et al, (2010) and Lauber et al. (2009) identified the same dominant groups, with the exception of Verrucomicrobia which was replaced by Nitrospira (Faoro et al., 2010) and TM7 and Cyanobacteria replacing Planctomycetes and Chloroflexi (Lauber et al., 2009).

### 4.2 Location, soil properties, climate and vineyard management contribute are associated with soil microbial community dissimilarity in the Barossa

Although dominant taxa were constant at a regional level, soil microbiome diversity and composition seemed to be a better factor separating soil microbiomes from different sub-regions. The phylogenetic inference of microbiome composition differences between sub-regions showed that OTU richer sub-regions (Northern and Central Grounds) clustered independently from the OTU poorer ones (Eden Valley and Western Ridge).

Previous studies have shown that the major factors determining compositional dissimilarities of soil microbiome between sites are dispersal constraints (which predicts that more distant soils should have greater phylogenetic dissimilarity) and environmental heterogeneity (Fierer, 2008; Liu et al., 2014; Burns et al., 2015). Analysis of the influence of geographical distance on soil microbiome composition differences between Barossa Valley Region vineyards showed a small significant correlation between both parameters. It could be argued that such small contributions to vineyard soil microbiome composition differences could be associated with the relatively small distances between the vineyards in this study (Average distance 11.7 km, minimum distance 0.7 km and maximum distance 26.5 km). However, this correlation was similar to that observed by Burns et al. (2015) when studying 19 vineyards of the Napa Valley AVA that were separated by up to 53 km. This suggests that dispersal constraints contribute to soil microbiome differences at a much smaller scale than previously perceived.

Environmental heterogeneity has been found to be more important than geographic distance in shaping bacterial community at different geographical scales (Fierer and Jackson, 2006; Miura et al., 2017; da C Jesus et al., 2009; Ranjard et al., 2013; Hermans et al., 2017). The main contributors to environmental associated variability in soil communities are differences in climatic conditions, topography, soil properties, and cultivation practices (Mezzasalma et al., 2018; Burns et al., 2015). Microbiome composition similarity analysis results did not show a clear clustering of vineyards according to their geographic location, indicating that even at a close geographic distance, environmental heterogeneity is the dominant factor shaping soil microbiome composition.

To determine which environmental factors contribute to the observed differences in soil microbial communities we used an automatic model building approach. This analysis revealed that when taken in combination, plant-available phosphorous and soil texture were the major contributors to soil microbiome differences between vineyards (approximately 20% of the total observed variability). Differences in plant-available phosphorous have been previously shown to impact microbial communities (Awasthi et al., 2011; Fierer and Jackson, 2006). In the studied vineyards, clay soils tended to show higher plant-available phosphorous content (Figure 4A & C), which is consistent with previous findings (Krogstad et al., 2005). Interestingly, soil particle size has been negatively correlated with microbiome community alpha diversity (Sessitsch et al., 2001) indicating that both variables could be affecting microbiome composition in an, at least partially, independent manner. Moreover, while genera *Streptomyces, Rubrobacter* (both Actinobacteria) and *unclassified MND1*, were especially prevalent in clay soils, genera *Streptomyces, Pseudomonas* and *unclassified Sinobacteraceae* were found in soils with plant-available phosphorous content higher than 30μg / g soil. *Pseudomonas*, are inorganic P solubilizing bacteria (Awasthi et al., 2011; Goswami et al., 2013; Schmalenberger and Fox, 2016). Conversely, P levels negatively correlated with the abundance of the organic P mineralizing taxon OPB35. Pairwise analysis of individual taxa and environmental variables also identified previously reported strong and positive correlations between soil pH and order iii1-15 (acidobacteria-6) and family Pirellulaceae (Rousk et al., 2010; Hermans et al., 2017; Wu et al., 2017). Interestingly, both P and pH, have been shown to be soil variables that are indicative of anthropogenic activity (Hermans et al., 2017), which highlights the potential use of such taxa as reliable indicators of soil condition.

Previous studies have shown that climatic variables such as rainfall (Wildman, 2015) and temperature (Cong et al., 2015) are major shapers of soil microbial population composition and activity. Our results indicate that cooler and wetter regions (Western Ridge and Eden Valley) had relatively lower soil microbial diversity, and a higher ratio of dominant species, than the warmer and drier sites. Additionally, elevation, which negatively affects air temperature, showed a positive correlation with the families Isosphaeraceae and an unsurprising negative correlation with the thermophilic taxon Conexibacteraceae (Wagner and Wiegel, 2008).

Agricultural lands tend to show similar patterns of dominant bacteria (Burns et al. 2015; Liu et al. 2014; Faoro et al. 2010; Lauber et al. 2009), indicating that microbial community composition can be profoundly affected by cropping practices (Hartman et al., 2018). Our results show that, both spacing between row and vine, which determine the vineyard’s planting density, are significantly associated with global differences in soil microbial community. Work in oil palm plantations has shown that planting density affects soil bacteria by altering the level of solar light incidence on soils, which can have dramatic effects on soil temperature and moisture (Tripathi et al., 2016). Pairwise comparisons between agronomical practices and individual taxa showed a negative correlation between spacing among vines on the same row and the abundance of representatives of the Haliangiaceae family. These are mesophilic organisms previously identified to be sensitive to agricultural practices (e.g. (Ding et al., 2014; Kim and Liesack, 2015; Wang et al., 2016)), which abundance could be favored by lower soil temperatures in densely planted vineyards. This highlights the importance of temperature, shown above, in the formation of soil bacterial communities. However, vine density and the use of under-vine cover crops could also cause different levels of interactions between plant roots and soil microbes. This is particularly prominent when comparing sites with similar topography and soil texture, in which spatial patterns of soil biota are assumed to be structured primarily by plant growth, age, growth form and density (Ettema, 2002). Our results indicate that the abundance of taxa from the bacterial family Hyphomicrobiaceae is positively correlated with the vineyard age. Plant age has previously been linked to differences in soil bacterial communities in annual crops (Marques et al., 2014; Walters et al., 2018) and in wild plant species (Wagner et al., 2016; Na et al., 2017). However, how composition and diversity of rhizosphere communities shift with plant age in perennial, long-living crops has received less attention. No-till soil management has been shown to affect community composition (Lewis et al., 2018). It is therefore tempting to speculate that in perennial crops, the effect of plant age on soil bacterial communities, is the result of the prolonged presence of the crop.

### 4.3 Correlations between soil bacterial communities and berry and wine parameters

Berry parameters were found to be significantly associated with both the composition and diversity of soil microbiomes and with the abundance of single taxa. A total of six fruit traits correlated with differences in bacterial community composition and diversity, while one fruit trait was found significantly associated with the abundance of specific taxa. Plant–microbe interactions are known to modify the metabolome of *Arabidopsis thaliana* plants grown under controlled conditions (Badri et al., 2013), however, the modulating effect of soil microbiomes on the metabolome of commercial crops is unexplored. Unfortunately, the non-intervention nature of this research impedes us determining if the relationships observed between vineyard soil microbiomes and fruit traits are causal or simply mere correlations.

Soil microbiomes have previously been described as a contributor to the final sensory properties of wines by affecting wine fermentation. Grape must microbiota was found to be correlated to regional metabolite profiles and was suggested to be potential predictor for the abundance of wine metabolites (Bokulich et al., 2016). Here we identified 19 wine traits correlated with differences in bacterial community composition and diversity, and seven correlated with the abundance of specific taxa. Vineyard soils may serve as a bacterial reservoir since bacterial communities associated with leaves, flowers, and grapes share a greater proportion of taxa with soil communities than with each other (Zarraonaindia et al., 2015). Unfortunately, the non-intervention nature of this research, the lack of replicability and the use of commercially produced wines, preclude us from determining if the relationships observed between vineyard soil microbiomes and fruit/wine traits are causal or simply mere correlations. Each of these wines was made commercially by different producers so there is potential for a certain level of winemaking effect.

## 5 Conclusion

Taken collectively our results show that geographic separation between vineyards contributes to bacterial community dissimilarities at a much smaller scale than previously reported. Environmental variables (e.g. climatic, topography, soil properties, and management practices) were the greatest contributor to such differences. Particularly, we found that soil variables are the major shapers of bacterial communities. Also, we show that variables highly affected by soil anthropogenisation (pH, plant available Phosphorous) and agricultural management variables (plant age, planting density) have strong correlations both with the community composition and diversity and the relative abundance of individual taxa. Finally, our results provide an important starting point for future studies investigating the potential influence of microbial communities on the metabolome of grapevines in general, and on the definition of local Terroirs. It will also be important to study a wider range of soil physicochemical properties, and vineyard floor vegetation, on the soil microbiome.

## 7 Author Contributions

TRC, JRS, AM, MG, JB, CC and CMRL conceived and planned the experiments. CC and RDB contributed to the design of the research project, vineyard selection and fruit and wine chemical characterization. JZ conducted soil physicochemical analysis and the 16S rRNA gene laboratory work. JZ and TMN conducted the bioinformatics analysis. JZ and CMRL took the lead in writing the manuscript. All authors provided critical feedback and helped shape the research, analysis and manuscript.

## 9 Acknowledgments

The authors would like to gratefully acknowledge the Barossa Grounds Project and in particular the growers that allowed us to sample material from their properties and supplied information about their vineyards and management strategies. Dr. Kendall R. Corbin contributed to soil sample collection. We are thankful to Dr Hien To, Dr Steve Pederson and Dr Rick Tearle for assistance with data analysis.

## 10 Data Availability Statement

All the data and supporting information will be made available online.

1. Supplementary Figures S1–S3.
2. Supplementary Tables S1-S11.

The data that support the findings of this study are openly available in NCBI Sequence Read Archive (accession number: PRJNA601984).

---

**Supplementary Figure 1.**
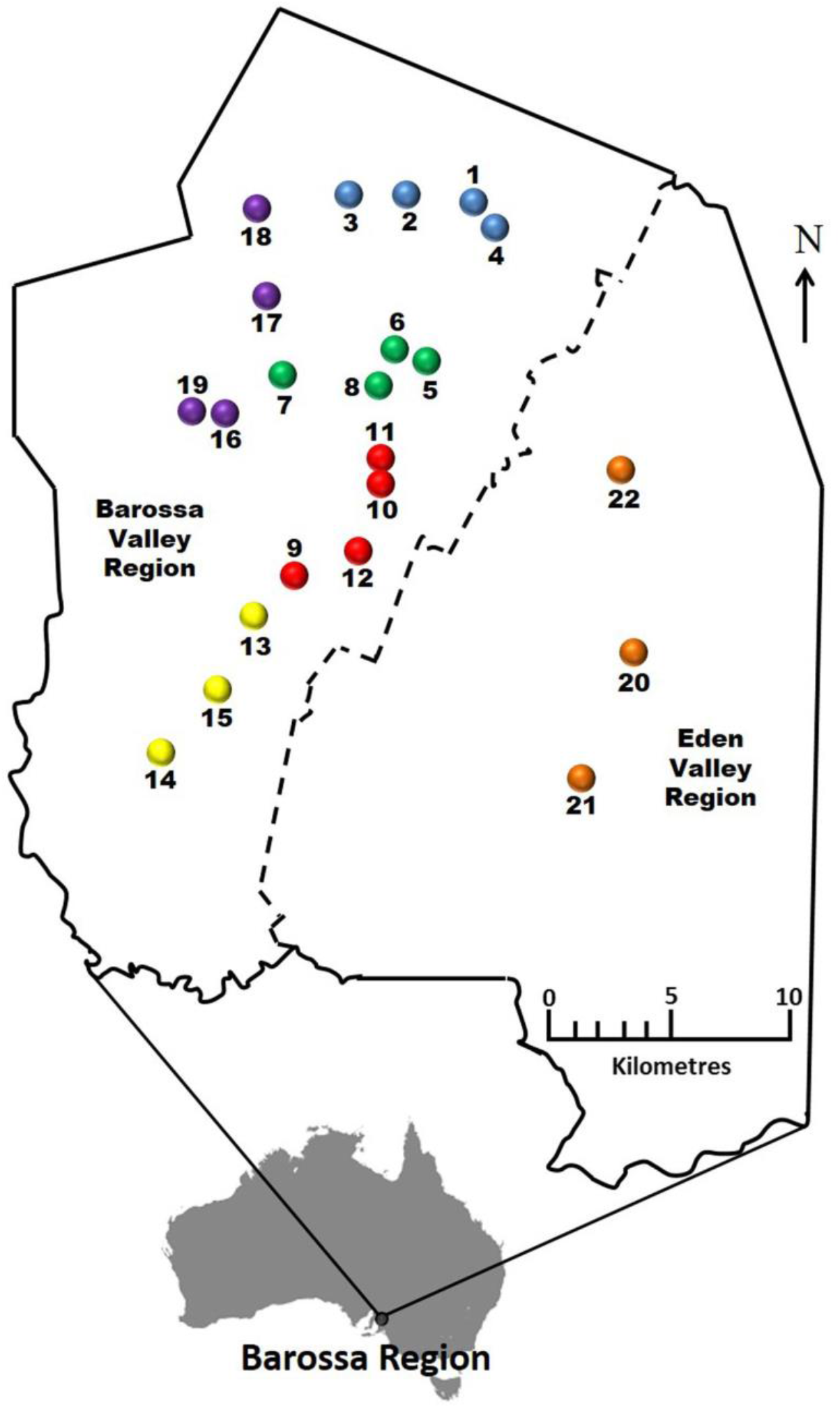
Location of 22 Barossa vineyard sites. Vineyards are color coded according to the six wine sub-regions as defined in Xie et al. (2017): Northern Grounds: Blue, Southern Grounds: Yellow, Central Grounds: Green, Eastern Ridge: Red, Western Ridge: Purple, Eden Valley: Orange. Map modified from Xie et al. (2017).

**Supplementary Figure 2.**
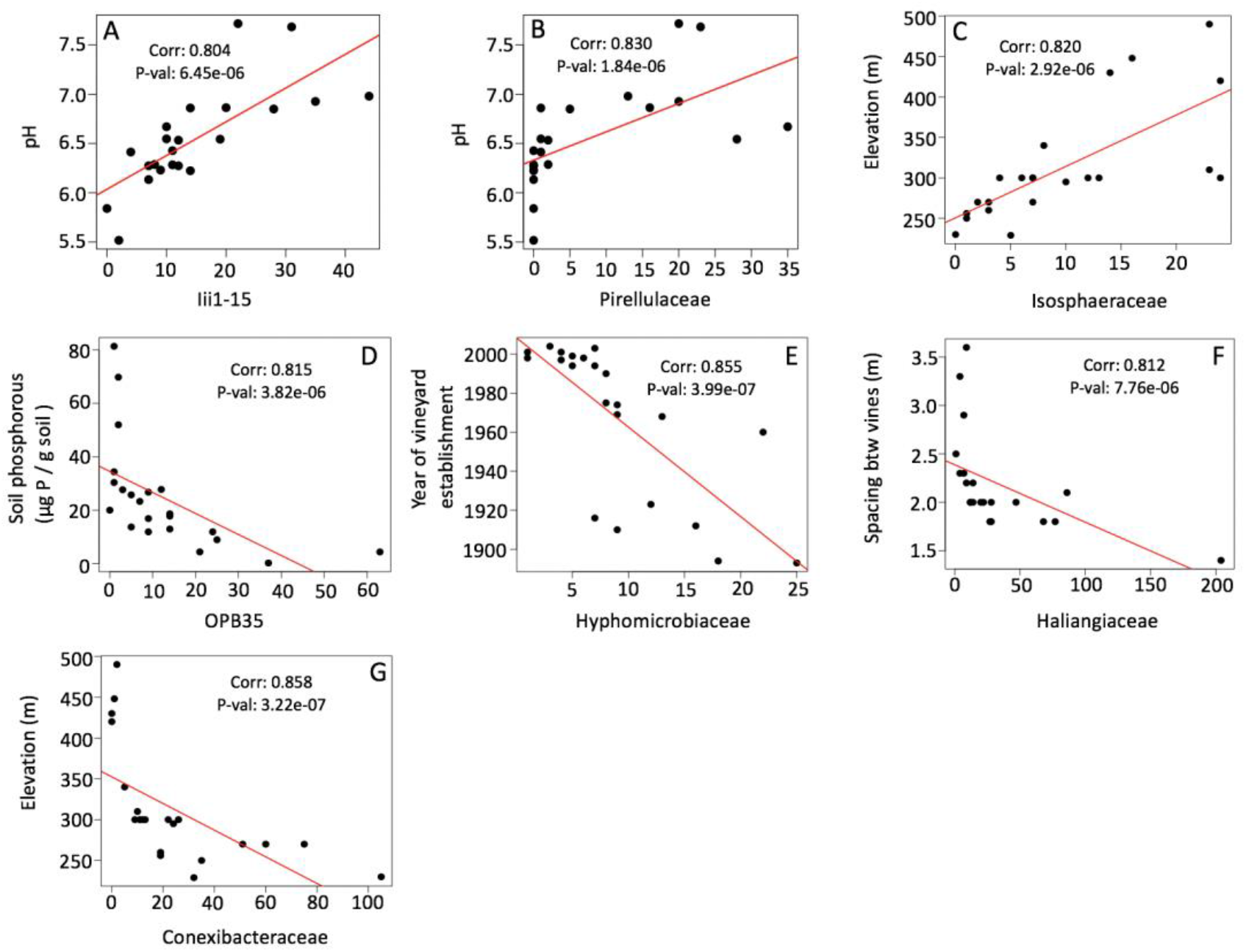
Association between taxon abundance and environmental/agronomical variables in Barossa Region vineyard soil bacteria communities. Correlations were tested using Spearman’s rank correlation coefficient with its probability estimate for significance (P) and implemented using the function *rcorr* in the R package *Hmisc*. Correlation coefficient and P values for each of the comparisons are included in each inset.

**Supplementary Figure 3.**
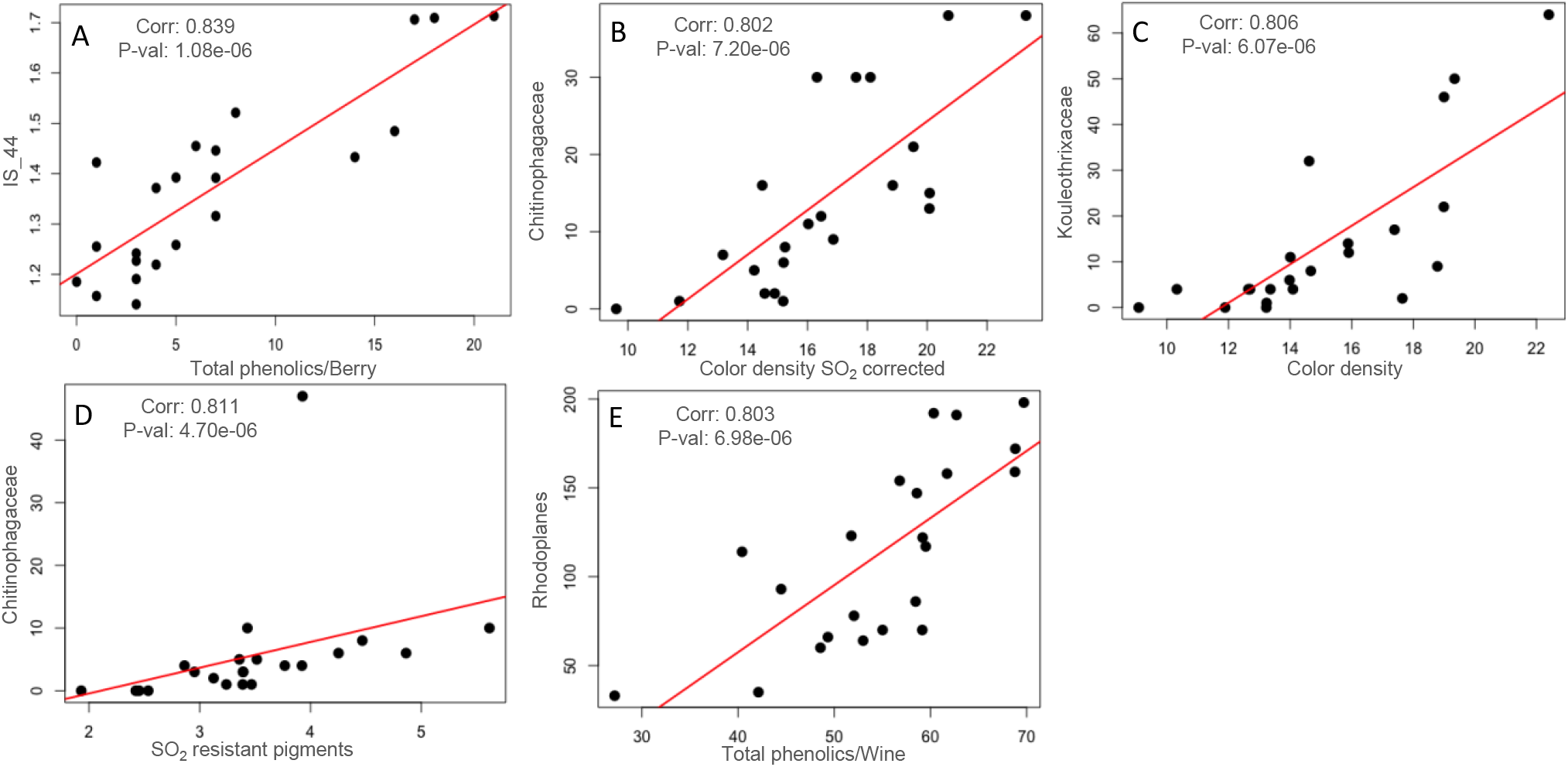
Association between taxon abundance and fruit/wine traits in Barossa Region vineyard soil bacteria communities. Relationship between taxon abundance and fruit (A) and wine (B-F) traits. Correlations were tested using Spearman’s rank correlation coefficient with its probability estimate for significance (P) and implemented using the function *rcorr* in the R package *Hmisc*. Correlation coefficient and P values for each of the comparisons are included in each inset.

